# Patient-specific zebrafish models reveal a complex phenotypic spectrum in *NBAS*-associated Atypical Osteogenesis Imperfecta

**DOI:** 10.64898/2026.07.24.740546

**Authors:** Daniel A. Baird, Sookyung Seo, Zuzanna Matelowska, Karthick Navin Mohandass, Alagumeenal Subramanian Annamalai, Aynour Abouelkhair, Myar Zafar, Nurhaziqah Supari, Sarah Baxendale, Catherine A. Loynes, Freek van Eeden, Meena Balasubramanian

## Abstract

Neuroblastoma amplified sequence gene (*NBAS*) variants are associated with short stature, optic atrophy, and Pelger-Huët anomaly (SOPH) syndrome. We previously identified compound heterozygous variants in *NBAS* to cause atypical Osteogenesis Imperfecta (OI), with these patients presenting with short stature, developmental delay and recurrent long-bone fractures. However, skeletal disease progression due to these variants and the disease mechanisms underlying *NBAS*-associated OI remain poorly understood. Here, we provide a clinical update on previously identified patients and investigate the role of NBAS during skeletal development using zebrafish knockout and patient-specific missense variant zebrafish models. Homozygous knockout larvae exhibited delayed operculum development, reduced bone ossification, and defects in Meckel’s cartilage morphology and its underlying cellular structure. Homozygous missense larvae displayed milder cartilage defects without any major defects to early skeletal structures. Seemingly opposing phenotypes were observed in compound heterozygous zebrafish carrying the knockout and missense alleles *in trans*, with no obvious phenotypes seen in the Meckel’s cartilage and accelerated operculum development observed. Together, our results demonstrate that different *nbas* variants differentially affect skeletal development, suggesting complex spectrums of phenotypic and pathogenic mechanisms in *NBAS*-associated atypical OI.

## INTRODUCTION

Skeletal structures are fundamental in providing a rigid framework for muscles and skin, protecting vital organs for life, and supporting efficient mineral exchange for homeostasis^1^. A primary component of bones is collagen, which has essential functions in maintaining skeletal structural integrity and elasticity for skeletal mechanical properties^2^. Due to the high abundance of collagen in the human body, mutations in genes related to collagen structure and/or function can have a significant impact on skeletal development and have a profound impact on patient morbidity^3–5^.

Individuals with structural or functional abnormalities to collagen can present with Osteogenesis Imperfecta (OI). OI patients are primarily characterised by increased susceptibility to fractures and decreased bone density^6, 7^. Two genes commonly disrupted in OI patients are *COL1A1* and *COL1A2*, with variants in these genes accounting for approximately 90% of OI cases^8^. The remaining cases are due to a variety of complex aetiologies and variants in non-collagen encoding genes, such as disruption to osteoblast maturation, deficiency of the collagen prolyl 3-hydroxylation complex, and protein chaperone defects^9^. It is believed that variants associated with OI lead to reduced collagen synthesis and secretion^10–12^. Despite this, knowledge on the disease progression and underlying mechanisms of these OI cases are not well understood.

We previously described two unrelated boys with atypical OI caused by compound heterozygous *NBAS* variants, carrying p.Arg1914His in trans with truncating variants p.Arg1004Ter or p.Gln678Ter. Both showed proportionate short stature, recurrent long-bone fractures, developmental delay, immunodeficiency and abnormal liver function tests. Fibroblasts from these patients had reduced NBAS protein expression and abnormal type I collagen morphology, suggesting a role for NBAS in collagen biology^13^. The p.Arg1914His is a C-terminal founder variant in the Yakut population and underlies SOPH syndrome when homozygous, characterised by short stature, optic atrophy and Pelger–Huët anomaly^14, 15^. Larger international cohorts have shown that missense variants in the Sec39 domain are typically associated with infantile liver failure syndrome type 2 (ILFS2), C-terminal variants such as p.Arg1914His with a predominantly multisystem SOPH-like phenotype, and β-propeller variants with a combined, severe hepatic and multisystem presentation^16, 17^. Within these series, p.Arg1914His is the single most frequent *NBAS* genotype and often occurs *in trans* with a truncating second variant, and the position of the missense or in-frame “determining” variant within the protein has emerged as the main correlate of clinical subtype^16, 17^. It is therefore important to define how the p.Arg1914His founder variant and loss-of-function *NBAS* variants disrupt protein function at the organismal level.

In this study, we have generated zebrafish models to investigate the disease-associated p.Arg1914His *NBAS* variant and null variants *in vivo*. By performing detailed phenotypic analyses, we have found structural and developmental abnormalities to developing collagen and skeletal structures in both models, with a phenotypic difference observed in compound heterozygous zebrafish models. By modelling *NBAS* variants in zebrafish, we have expanded on the current understanding of how NBAS dysfunction impacts skeletal and collagen development *in vivo*.

## METHODS

### Clinical data collection and literature review

Electronic medical records were reviewed to obtain updated phenotypic information of our patients with *NBAS* compound heterozygous variants we previously reported^13^. Clinical data included detailed phenotypic assessments extracted using a structured proforma. Written informed consent for publication of clinical data was obtained from both families. A narrative literature review was conducted in PubMed and Google Scholar from inception until June 2026, to obtain an update in extant literature to understand phenotypic evolution of NBAS-related phenotype.

### Amino acid and protein conservation analysis

Amino acid similarities between NBAS proteins in human (Uniprot accession number: A2RRP1), mouse (Uniprot accession number: E9Q411), and zebrafish (Uniprot accession number: Q5TYW4) were compared using Protein Blast software^18^. Analysis of the conservation and amino acid similarity of the specific p.Arg1914His variant was performed using the Clustal Omega software^19^.

### Zebrafish husbandry

Zebrafish strains were raised and housed at the Bateson Centre for Disease Mechanisms at the University of Sheffield. Aquaria used to house zebrafish are approved by the UK Home Office and maintained in accordance with the UK Animals (Scientific Procedures) Act 1986. Wildtype strains were from stocks held at the University of Sheffield Biological Services Unit aquaria. Zebrafish were raised in a constant 14:10 hour light-dark environment at 27-28°C according to standard protocols.

Maintenance and staging of larvae were performed as described previously^20^. All experiments were conducted according to the project licences (PP5148348 and PP3627554) approved by the UK Home Office.

### Zebrafish line generation & maintenance

Upon confirmation of homology at the patient variant amino acid site, a p.Arg1919His (Arg1919His) mutation was generated at the University of Sheffield, with this line being termed *nbas^sh680^*. The zebrafish *nbas* gene (ENDSDARG00000008583) was targeted with CRISPR-Cas9 guides, enzymes and a homology directed template (Supplementary Table 1) by injections at the one-cell stage. F0 embryos were raised to adulthood and outcrossed to an AB wildtype strain, where germline transmission was confirmed in the F1 generation. F1 generation mutants were then outcrossed to either AB wildtype, *Tg(sp7:GFP; col2a1:mCherry*), or *Tg(sox10:GFP; col2a1:mCherry)* lines. Sanger sequencing to confirm DNA edits was performed with primers listed in Supplementary Table 1. Impact on protein sequences were predicted using the ExPASy software^21^ and Vector Bee (https://www.vectorbee.com/).

All mutant analyses were performed on F3-F5 larvae generated by inbreeding of heterozygous adult fish. As controls, a mix of either +/+ and +/- siblings from a +/- adult inbreeding or +/+ larvae from +/+ adult inbreeding (siblings of +/- adults) were used. Genotyping was performed by PCR with primers shown in Supplementary Table 1. Following PCR, DNA products underwent restriction digestion with either SpeI-HF (#R3133S) or AvaI (#R0152S) for *nbas^sa16290^* and *nbas^sh680^* lines respectively.

### Transgenic imaging and operculum image analysis

Larvae were selected based on their transgenic status for mutants in either the *Tg(sp7:GFP; col2a1:mCherry)* or *Tg(sox10:GFP; col2a1:mCherry)* backgrounds. Larvae negative for transgene expression were used for either Alizarin red or calcein stainings. Larvae with positive transgene expression were anaesthetised with MS-222 and mounted in either the lateral or ventral orientation. Images with the desired fluorescent filters with either a Zeiss AxioZoom v16 fluorescent stereoscope or a Nikon W1 spinning disk confocal microscope were taken. Brightfield images of larval heads were also taken on a Zeiss AxioZoom v16 fluorescent stereoscope. The length and width of the operculum was measured from transgenic samples alongside length and width measurements of the head from brightfield images for normalisation. Ratios of the operculum to head dimensions were then calculated.

### Alcian blue cartilage staining

Cartilage staining using Alcian blue was performed as previously described with minor modifications^22, 23^. Larvae were fixated overnight in 4% PFA at 4°C. They were then washed with PBST, bleached (3% H_2_O_2_, 2% KOH in ddH_2_O) for 1 hour, and then washed thrice in PBST. Samples were stained with 0.01% Alcian blue/60 mM MgCl_2_/70% EtOH overnight at 4°C. After overnight incubation, larvae were incubated for 10 minutes at room temperature successively with 80% EtOH/10 mM MgCl_2_, 50% EtOH/10 mM MgCl_2_, and 25% EtOH/MgCl_2_. Additional bleaching was performed for 30 minutes before larvae were washed a minimum of two times with 25% glycerol/0.1% KOH. Samples were stored in 50% glycerol/0.1% KOH at 4°C before imaging. Larvae were imaged in the ventral orientation using either a Nikon SMZ18 stereoscope. Distances and angles of cartilage were measured as previously described^22^.

### Fixed Alizarin red staining

Skeletal staining with fixed Alizarin red staining was performed as previously described^24^. Briefly, larvae at 5 dpf were euthanised and fixated (5% formalin, 5% Triton X-100, 1% KOH in PBS) at 42°C for a minimum of 3 days. Samples were then washed in 20% ethylene glycol/1% KOH for 5 minutes before being stained with 0.05% Alizarin red S/20% ethylene glycol.1% KOH or 30 minutes at room temperature. Samples were placed in clearing solution (20% Tween20/1% KOH) at 42°C for 30 minutes before a glycerol series was performed. Samples were then mounted and imaged with a Zeiss AxioZoom v16 fluorescent stereoscope. Images were analysed using Fiji/ImageJ software^25^.

### Calcein green live staining

Live staining was performed as previously described^26^. In brief, larvae were submerged in either 0.2% calcein green (pH 7.4) in the dark at 28°C for 30 minutes maximum. Larvae were then washed 3 times in clear E3 embryonic media for 5 minutes each time, before larvae were immediately mounted.

Fluorescent images were taken on a Zeiss AxioZoom v16 fluorescent stereoscope. Images of the operculum had their areas measured and integrated density calculated using Fiji/ImageJ software^25^.

### Data Normalisation and Statistical Analysis

Previous studies have noted a high degree of variability in the staining and some skeletal/cartilage morphological features detected by Alcian blue, Alizarin red, and calcein stainings. Significant variability was seen between different clutches of embryos born at different times and within the same clutch born at the same time^27^. Skeletal and cartilage developmental timings can differ between individual embryos, making some changes in structures, mineralisation and calcification difficult to detect Differences in these parameters between experiments, and within, could mask subtle abnormalities and differences between genotypes. To account for inter- and intra-clutch variability, data was normalised for all independent experiments to either the +/+ or sibling embryos/larvae for each independent repeat with the exception of operculum ratio measurements.

Data are shown as a boxplot, with individual datapoints from experiments overlaying the plot. Representative images and graphs are presented. All results from experimental protocols were validated by independent repetitions carried out n≥ 3 times. Data was tested for Gaussian distribution by a Shapiro-Wilk test. Statistical differences and variance between groups were tested by either student’s t-test, ordinary one-way analysis of variance or Kruskal-Wallis test. Significance levels are as follows: * *p* < 0.05, ** *p* < 0.01, *** *p* < 0.001, ns: not significant.

## RESULTS

### Updated clinical phenotype of two patients with p.Arg1914His

Across published cohorts, more than 200 individuals with biallelic NBAS variants have been described, predominantly with hepatic, growth, ophthalmological, immune and skeletal manifestations. In addition to our two patients, a few other case reports have documented bone fragility as a feature. Our current work adds longitudinal follow-up of two patients with p.Arg1914His in trans with a truncating NBAS variant.

Our patient 1 is now 21 years old and has successfully transitioned to adult services. Since the last report at age 10 years, his phenotype has evolved to include mild, infection-triggered transaminase elevation that has lessened with age, with investigations demonstrating oesophageal varices and portal hypertensive gastropathy; he underwent percutaneous gastrostomy closure at 14 years. He developed insulin-dependent diabetes mellitus at 16 years and was subsequently diagnosed with primary testicular failure. He continues to receive 6-monthly zoledronic acid for low bone mineral density, with good skeletal response, and his growth remains below the 0.4th centile. Immunoglobulin replacement was maintained for recurrent viral infections; despite cytopenias and splenomegaly, there was no evidence of bone marrow failure, supporting peripheral consumption as the likely mechanism. He is now registered blind with history bilateral optic atrophy and myopia. He also has a specific learning disability and autism yet continues to attend college.

Patient 2 is now 16 years old and has transitioned from childhood into ongoing multidisciplinary follow-up. Since our last report at age 6 years, he has sustained further fractures, including tibial diaphyseal, clavicular, and femoral fractures, the latter requiring surgical correction, with consequent reduction in ambulation and the need for a walking frame and wheelchair. He has had a clear response to pamidronate, with improved bone mineral density, and continues 3-monthly treatment. Growth remains below the 0.4th centile, with associated mild learning difficulties. His hepatic phenotype is recurrent infection-triggered acute liver failure, which has become less frequent with age. He also has hypogammaglobulinemia requiring immunoglobulin replacement and no evidence of bone marrow failure has been reported. Endocrine and other systemic features include glutamic acid decarboxylase antibody-positive type 1 diabetes mellitus diagnosed at 9 years, mild aortic regurgitation on cardiac imaging, bilateral optic atrophy, cone dysfunction, myopia, and unilateral exotropia, with preserved hearing.

### Amino acid conservation between species allows for *NBAS* missense model generation in zebrafish

To assess whether we could generate a zebrafish model for the p.Arg1914His founder variant, we performed Protein Blast and multiple sequence alignment of human, mouse and zebrafish protein sequences to ensure that the arginine amino acid at position 1914 in the human protein was conserved between species. Protein blast analysis revealed that the zebrafish Nbas protein was 63.85% identical to the human protein, and 62.26% identical to the mouse protein. Conservation of the arginine was found between human and zebrafish, with this specific arginine amino acid in zebrafish found at position 1919 in the protein sequence (Fig. 1A). This allowed us to generate a knock-in zebrafish model (*nbas^sh680/sh680^*) of the human p.Arg1914His missense variant (p.Arg1919His in zebrafish) using CRISPR-Cas9 and homology-directed repair (Fig. 1Bi). Protein sequence prediction showed that the nucleotide changes at this site does result in the wildtype arginine being substituted for a histidine (Suppl. Figs. 1i & ii). This p.Arg1919His zebrafish model will now be referred to as the *nbas* ‘missense’ model. Furthermore, we obtained a zebrafish knockout (KO) model of *nbas* with a c.4316G>A point mutation in exon 37 (*nbas^sa16290/sa16290^*) confirmed with Sanger sequencing (Fig. 1Bii), with protein sequence analysing predicting that this point mutation results in a premature stop codon being introduced just after a predicted Sec39 domain (Suppl. Fig. 1iii). This null mutation zebrafish line will now be referred to as the *nbas* ‘KO’ model. Both homozygous KO and missense *nbas* zebrafish models did not show overt skeletal phenotypes at 5 dpf (Fig. 1C).

**Figure 1.**
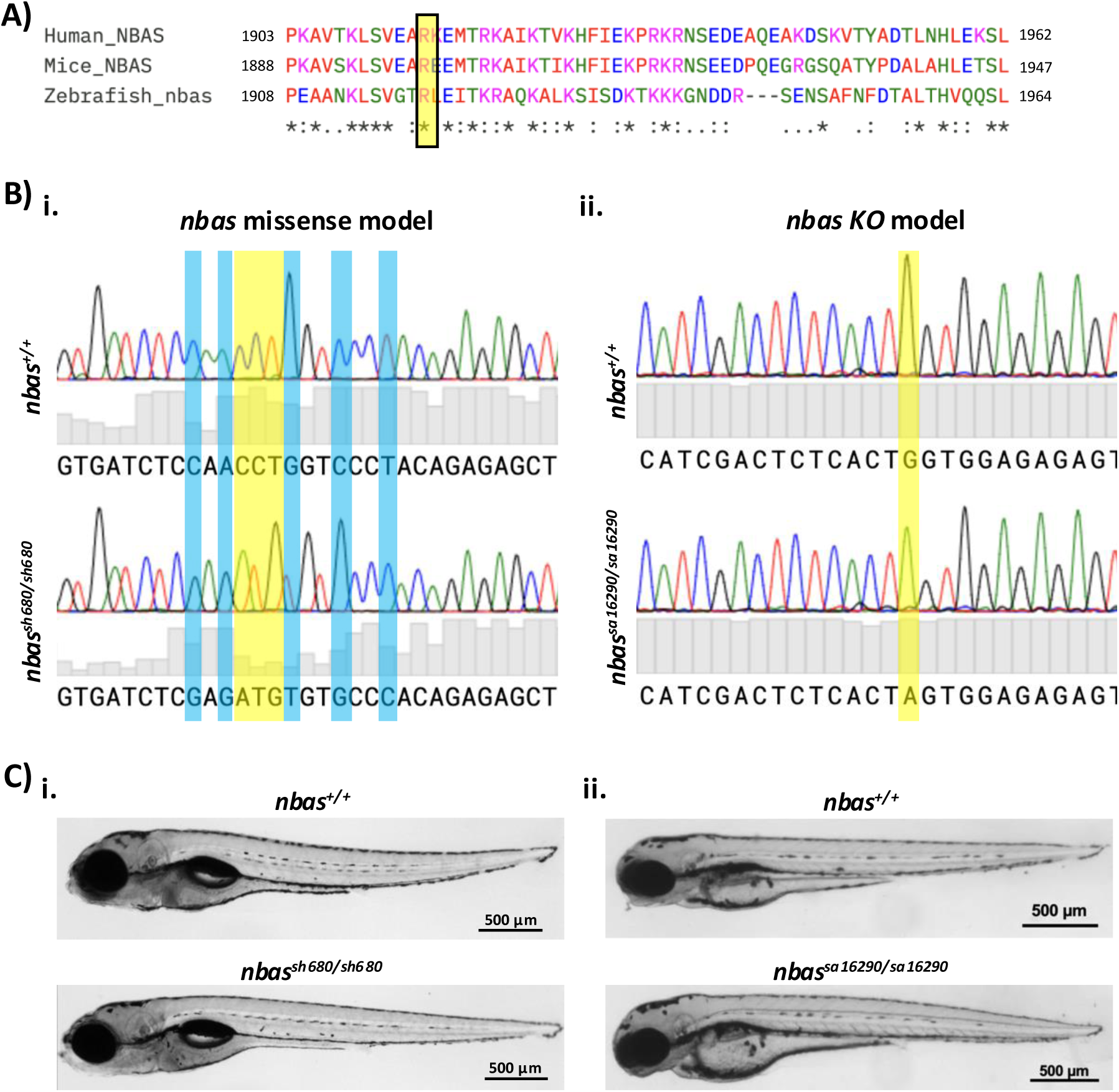
Generation of *nbas* zebrafish models to investigate patient-specific variants. **A.** Amino acid conservation analysis of neuroblastoma amplified sequence protein between humans, mice, and zebrafish using Clustal Omega software. Yellow box highlights patient p.Arg1914His variant. ‘*’ represents fully conserved amino acid, ‘:’ represents highly conserved amino acid, and ‘.’ represents weakly conserved amino acid. **B.** Sequencing chromatograms for zebrafish *nbas* models. **i)** Sequencing chromatogram for *nbas* sh680 allele missense model with wildtype on top panel and homozygote on the bottom. Highlighted yellow bar shows mutation to sequence to induce arginine to histidine substitution. Highlighted blue bars represent silent mutations. **ii)** Sequencing chromatogram for *nbas* sa16290 knockout model with wildtype on top and homozygote on bottom panel. Highlighted yellow bar shows mutation predicted to cause protein truncation. **C.** Gross morphology brightfield images of 5 dpf larvae. **i)** Wildtype larvae displayed on the top panel with *nbas^sh680/sh680^* missense homozygous larvae on the bottom. **ii)** Wildtype larvae displayed on the top panel with *nbas^sa16290/sa16290^* knockout homozygous larvae on the bottom. Scale bars, 0.5 mm.

### Morphological and ossification defects in early opercula structures found in *nbas* KO model, but not in missense model

To study the early impact of *nbas* variants on skeletal development, we investigated the structure of the operculum - one of the earliest skeletal structures to form in zebrafish. Ratios of operculum lengths and widths with the respective length and width of the head were calculated to ensure that inter-individual size differences did not affect analysis (Figs. 2Ai & ii). We found that both the length and width ratios of operculum were significantly smaller in *nbas* homozygous KO larvae when compared to wildtype and heterozygous siblings (will be referred to as “siblings” hereafter for both KO and missense models) from 3 dpf up to 5 dpf (Fig. 2C). However, no differences at any timepoint between 3 and 5 dpf were observed between the *nbas* homozygous missense larvae and their siblings for either length or width operculum ratios (Fig. 2D).

**Figure 2.**
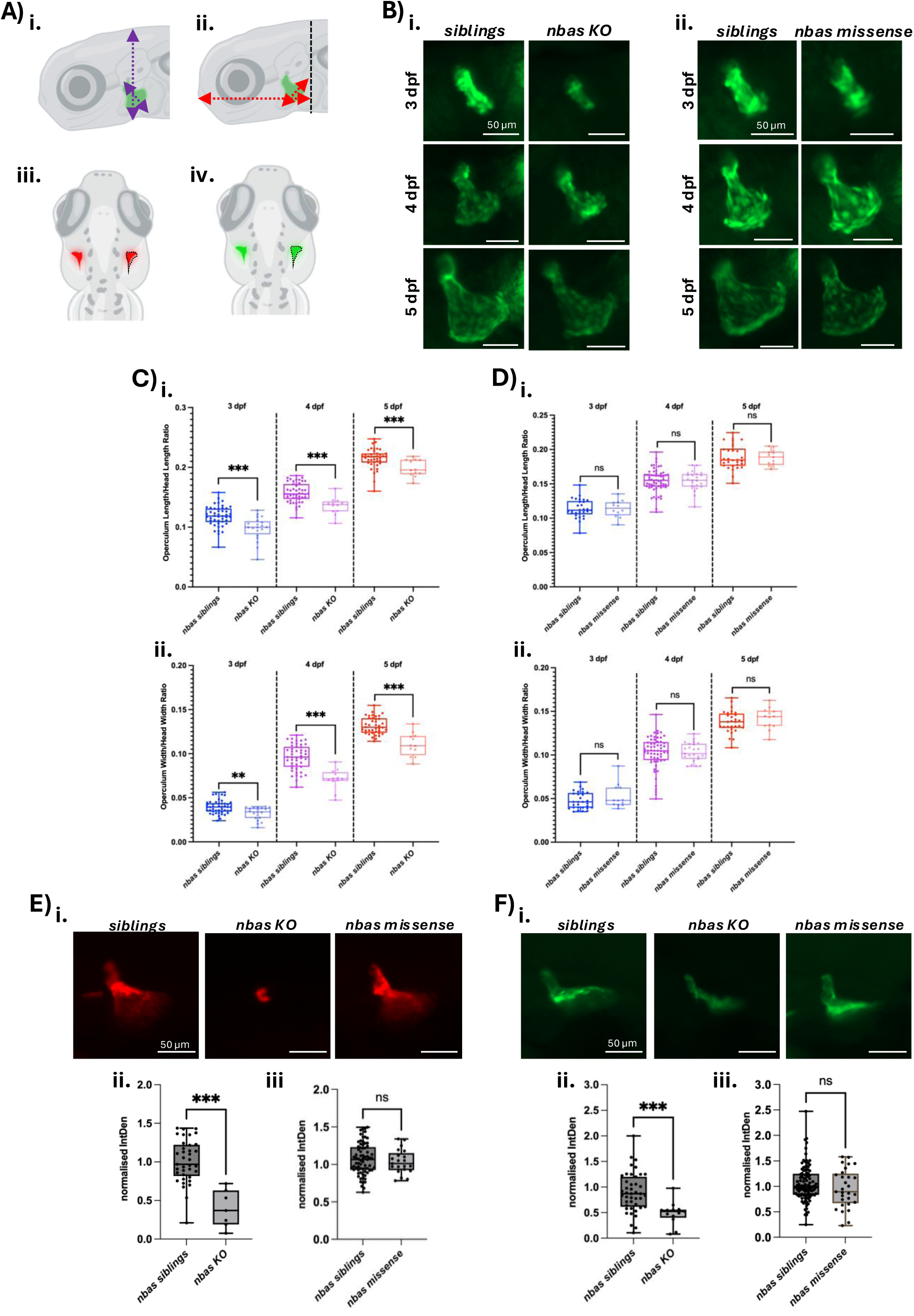
Operculum morphology and mineralisation disrupted in knockout larvae. **A.** Schematic showing how operculum **i)** length and **ii)** width were measured normalising to the length and width of the head. Integrated density of operculum measured from area shown in **iii)** for Alizarin red and **iv)** for calcein stainings. Schematic created with BioRender. **B.** Transgenic immunofluorescence of the operculum using *Tg(sp7:GFP)* for **i)** knockout and **ii)** missense models. Upper panels show operculum structure at 3 dpf, middle panels at 4 dpf and bottom panels at 5 dpf. Scale bars, 0.05 mm. **C.** Quantification of operculum morphology in *nbas* knockout models from 3 to 5 dpf for **i)** operculum length to head length ratio and **ii)** operculum width to head with ratio. Between each genotype group within each day, either an unpaired t test or a Mann-Whitney test was performed. **D.** Quantification of operculum morphology in *nbas* missense models from 3 to 5 dpf for **i)** operculum length to head length ratio and **ii)** operculum width to head with ratio. Between each genotype group within each day, either an unpaired t test or a Mann-Whitney test was performed. **E.** Alizarin red stainings. **I)** Alizarin red staining fluorescent images of the operculum at 5 dpf for representative siblings, knockout and missense models. Scale bars, 0.05 mm. Quantification of Alizarin red normalised integrated density for **ii)** knockout and **iii)** missense model. **F.** Calcein green stainings. **I)** Calcein green staining fluorescent images of the operculum at 5 dpf for representative siblings, knockout and missense models. Scale bars, 0.05 mm. Quantification of Alizarin red normalised integrated density for **ii)** knockout and **iii)** missense model. Ns, not significant. ** *p* < 0.002. *** *p* < 0.001.

To address potential osteogenic process differences in operculum development at 5 dpf, we performed Alizarin red and calcein stainings as outlined (Figs. 2Aiii & iv). In homozygous KO larvae, we observed a significant decrease in the normalised integrated density fluorescence intensity of the operculum in both stainings. Again, *nbas* missense homozygous larvae did not show any fluorescent differences when compared to sibling counterparts in either Alizarin red or calcein stains (Figs. 2E & F). Together, these results suggest that the KO mutation in *nbas* can cause morphological and osteogenic development abnormalities to the early operculum structure, whereas the missense model does not show these defects at this timepoint.

### Abnormal development of the Meckel’s cartilage observed in both *nbas* zebrafish models

We investigated the development of the cartilage in our *nbas* models to determine whether skeletal deformities could be arising due to cartilaginous abnormalities. Alcian blue stainings for cartilage in the craniofacial regions were performed at 5 dpf (Fig. 3A), with differences seen in the Meckel’s cartilage. We observed that the distance between the two extreme points of the Meckel’s cartilage (M-M distance) were significantly shorter in both homozygous KO and missense models compared to their sibling counterparts (Figs. 3Bi & Ci). However, no differences in the angle of the Meckel’s cartilage between the two extreme points (M angle) was found in either genotype or model (Figs. 3Bii & Cii). Notably, the posterolateral edges of the Meckel’s cartilage appeared thickened and curved inwards in *nbas* KO homozygotes. Thus, we measured the thickness of his cartilage and found that KO Meckel’s cartilage thickness was significantly thicker compared to siblings. This trend was not observed in the missense model at the same timepoint (Figs. 3Biii & Ciii). As well as morphological defects in Meckel’s cartilage, the inner perimeter length (arc length) traced from one posterior edge to the other presented a significant decrease in KO homozygotes *Tg(col2a1:mCherry)* (Suppl. Fig. 2A), alongside a reducing trend in Alcian blue staining analysis. No difference was found in missense homozygous larvae (Suppl. Fig. 2).

**Figure 3.**
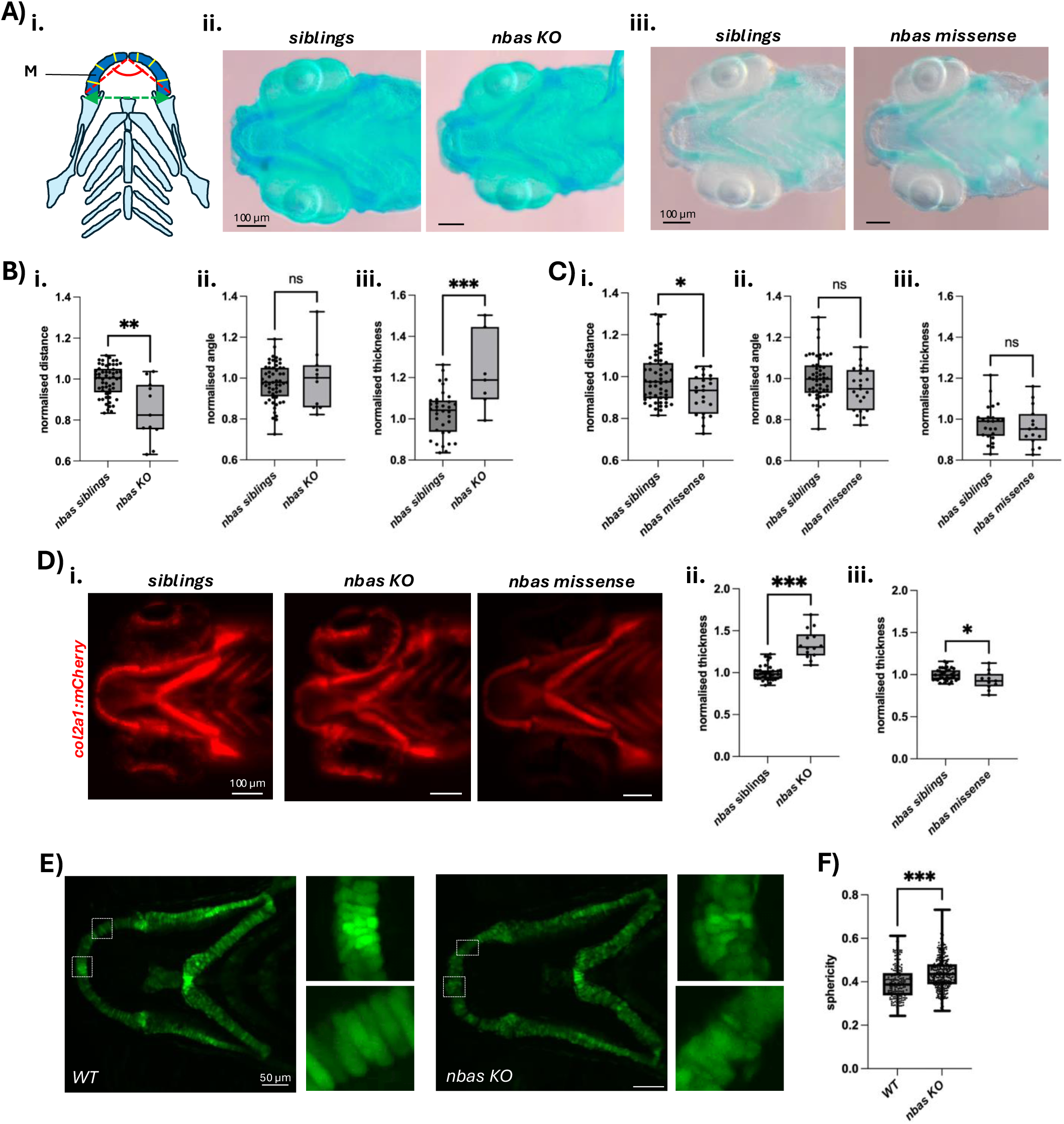
Meckel’s cartilage morphology defects in *nbas* knockout & missense models. **A.** Alcian blue stainings of craniofacial cartilage with **i)** schematic showing measurements taken for Meckel’s cartilage (M). Green arrow shows M-M distance. Red angle highlights M-M angle. Yellow lines show six measurements taken for each Meckel’s cartilage thickness which is then averaged. Representative images of Alcian blue stainings for **ii)** knockout and **iii)** missense models. **B.** Quantification of normalised Meckel’s cartilage measurements comparing knockout homozygous larvae and siblings at 5 dpf for **i)** distance, **ii)** angle, and **iii)** thickness. An unpaired t test or a Mann-Whitney test were performed. **C.** Quantification of normalised Meckel’s cartilage measurements comparing missense homozygous larvae and siblings at 5 dpf for **i)** distance, **ii)** angle, and **iii)** thickness. An unpaired t test was performed for statistical analyses. **D. i)** Representative immunofluorescent images of *Tg(col2a1:mCherry)* larvae at 5 dpf. Scale bars, 0.1 mm. Quantification of type II collagen layer normalised thickness between siblings and homozygotes for **ii)** knockout and **iii)** missense models. **E.** Representative immunofluorescent images of wildtype and homozygous knockout larvae at 5 dpf for *sox10:GFP*. White boxes highlight enlarged frames. Scale bars, 0.05 mm. **F.** Quantification of cell sphericity in Meckel’s cartilage between 5 dpf wildtype and homozygous knockout larvae. Ns, not significant. * *p* < 0.033. ** *p* < 0.002. *** *p* < 0.001.

Additionally, the angle between the ceratohyal (CH) cartilages showed a significantly wider trend in KO homozygotes compared to siblings, with a non-significant wider angle observed in between the Meckel’s and palatoquadrate (M-PQ) cartilages. No differences were found in either model between the PQ-PQ or M-CH distances. Interestingly, the thickness of the CH cartilage in KO homozygous mutants was significantly larger than siblings, but was found to be thinner in missense homozygotes (Suppl. Fig. 3).

### Variants in the *nbas* gene disrupt the underlying structure of the Meckel’s cartilage

Transgenic imaging for type II collagen revealed dysmorphic architecture of the Meckel’s cartilage accompanied by abnormal anterior/posterior medial curvature and widened mandibular symphysis in *nbas* KO homozygotes compared to siblings. No clear visible differences were observed between missense homozygous larvae and their sibling counterparts (Fig. 3Di). Furthermore, it was found that the layer of *col2a1* positive cells was significantly thicker in KO homozygotes but this layer was thinner in missense homozygous larvae, as is seen in the cartilage thickness (Figs. 3Dii & iii). The increased thickness of *col2a1* positive cells was also observed in the ceratohyal cartilage of KO homozygotes. No differences in the thickness of this layer was seen in the missense model.

We found that cells expressing *sox10* within the Meckel’s cartilage showed irregular geometry in *nbas* KO homozygous larvae at 5 dpf. Within wildtype larvae, *sox10* positive cells showed regular, stacked morphology, whereas these cells in KO homozygotes displayed increased sphericity and were patterned in a disordered manner (Fig. 3E & F).

### Milder developmental defects in operculum and Meckel’s cartilage observed in patient-specific compound heterozygous *nbas* model

Based on the evidence of cartilaginous and skeletal defects of both *nbas* knockout and missense models, we investigated a patient-specific zebrafish models (compound heterozygous) having missense and nonsense PTC variants on each allele through outcrossing heterozygous adults (Fig. 4A). We found that the length ratio of operculum was larger in *nbas* compound heterozygous larvae compared to wildtype, while width ratio was comparable to that of wildtype at 5 dpf (Fig. 4B).

**Figure 4.**
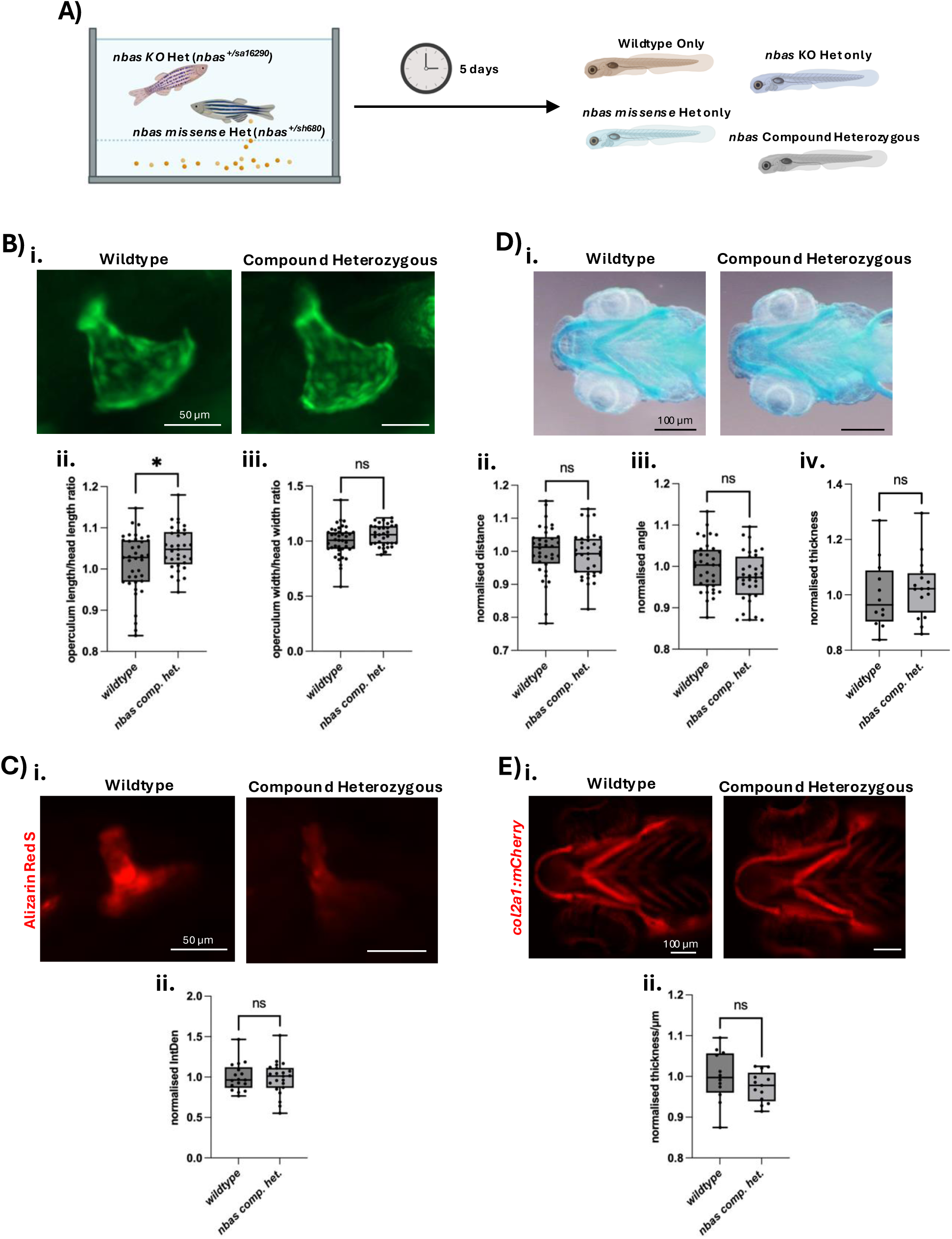
Operculum but neither cartilage or skeletal abnormalities found in patient-specific compound heterozygous *nbas* model. **A.** Schematic showing the generation of *nbas* compound heterozygous model through breeding of heterozygous *nbas* knockout and missense adult zebrafish. Created with BioRender. **B. i)** Representative images of opercula at 5 dpf from *Tg(sp7:GFP)* wildtype and compound heterozygous larvae. Scale bars, 0.05 mm. Quantification of **ii)** operculum length to head length and **iii)** operculum width to head width. Statistical analyses performed using a Mann-Whitney test. **C. i)** Representative images of Alizarin red stained opercula in 5 dpf wildtype and compound heterozygous larvae. Scale bars, 0.05 mm **ii)** Quantification of normalised integrated density of the operculum. An unpaired t test was used for statistical analysis. **D. i)** Representative Alcian blue images of craniofacial cartilage in 5 dpf wildtype and compound heterozygous larvae. Scale bars, 0.1 mm. Quantification of Meckel’s cartilage measurements between wildtypes and compound heterozygotes at 5 dpf for **ii)** distance, **iii)** angle, and **iv)** thickness. Unpaired t tests were used for statistical analyses. **E. i)** Representative images of *Tg(col2a1:mCherry)* in the craniofacial region of 5 dpf wildtype and compound heterozygous larvae. Scale bars, 0.1 mm. **ii)** Quantification of type II collagen layer normalised thickness. Unpaired t test used for analysis. ns, not significant. * *p* < 0.033.

However, the extent of calcification of the operculum and Meckel’s cartilage’s morphologies including distance, angle and thickness were found to be similar between patient-specific and wildtype larvae in Alizarin red and Alcian blue stainings (Figs. 4C & D). Interestingly, the arc length of Meckel’s cartilage of compound heterozygous larvae was increased compared to wildtype larvae (Suppl. Fig. 2B). In addition, the thickness in *col2a1* expressing cells in the Meckel’s cartilage’was similar between compound heterozygous variants and wildtype larvae, despite a decreasing trend that was not statistically significant being seen (Fig. 4E). No structural abnormalities or differences in *col2a1* positive cell layers in any of the other craniofacial cartilages were observed in the *nbas* compound heterozygous model (Suppl. Fig. 3).

## DISCUSSION

### Clinical follow-up of previously identified patients supports *NBAS* involvement in skeletal disorders

Variants in *NBAS* are associated with SOPH syndrome, infantile liver failure syndrome type 2, and atypical OI^13–17^. Here we provide long-term clinical follow-up of two patients with the C-terminal founder p.Arg1914His variant in trans with a truncating variant and interpret their phenotypic evolution in the context of expanding NBAS literature. Our observations reinforce three key points: existence of prominent bone fragility in SOPH-phenotype, variable hepatic involvement even within C-terminal genotype, and importance of multi-system surveillance from diagnosis into adulthood.

Our two patients have persistent severe short stature, recurrent low-impact fractures, reduced bone mineral density requiring bisphosphonates to stabilise them. Together with our previous original work showing reduced NBAS protein and abnormal type 1 collagen fibrils in patient-derived fibroblasts, these clinical findings support the role of NBAS in collagen biology and function within the skeletal system.

### Loss of *nbas* disrupts early skeletal development

To investigate how distinct *NBAS* variants influence protein function and development at the organismal level, zebrafish were utilised as a disease model due to their optical transparency, fast developmental timeframes, and high fecundity^28, 29^. Furthermore, zebrafish share a high degree of genetic conservation with humans, with 71% of human genes having an ortholog in zebrafish, and approximately 80% of genes associated with human disease have a corresponding zebrafish counterpart^30^. Sequence alignments demonstrated protein conservation of the affected arginine amino acid between zebrafish and humans, enabling zebrafish modelling of this specific missense variant. Zebrafish also have similar skeletal physiology and pathology^31, 32^ . Altogether, zebrafish are particularly suitable for investigating disease variants associated with OI and overall skeletal development.

Early skeletal development was assessed through analysis of the opercular bone, as this is one of the dermal first bones to form and ossify in zebrafish and is a well-established marker of osteogenic activity^33, 34^. We found that in *nbas* KO homozygous larvae, the opercular bone had pronounced dysmorphic features from 3 to 5 dpf, indicating skeletal development disruption at an early stage.

Growth of the operculum occurs at its fastest rate between 4-6 dpf^34^. As defects are observed at 3 dpf, it is possible that abnormalities in skeletal development arise before or during osteoblast differentiation. Opercular phenotypes we observe in our KO model contrast with previously characterised zebrafish OI models, in which the operculum is unaffected during comparable developmental stages^35^. Since operculum growth and mineralisation is driven by osteoblastogenesis, and that osteoclasts are not primarily present at this developmental timepoint, the abnormal structure of the operculum is likely reflective of impaired osteoblast function or differentiation^34, 36^.

Dysmorphic opercular structural features were accompanied by decreased mineralisation and calcification of the opercular bone in 5 dpf KO homozygotes. In human osteoblast cells, reduced mineral deposition is indicative of abnormal bone remodelling in osteoporosis^37^, which may result in increased bone fragility. A potential underlying mechanism involves matrix Gla protein (MGP), as this is believed to interact with NBAS to regulate bone mineralisation. Up-regulation of *MGP* gene expression following NBAS protein depletion has been shown to inhibit bone formation^13, 38^, which is consistent with the phenotype we observed in KO homozygous larvae. In contrast, no phenotypic differences in either the structure, mineralisation, or calcification of the operculum were observed in the *nbas* missense model between 3 to 5 dpf. This indicates that the missense and truncating *nbas* variants differentially affect Nbas function during early skeletal development.

### Variant-specific impacts on early cartilage development

Since craniofacial bones of zebrafish can initially develop from cartilaginous templates^39, 40^, craniofacial cartilage was examined to determine whether disrupted bone development and ossification may have a cartilaginous origin. Alcian blue staining revealed a reduction in the Meckel’s cartilage distance - the most anterior craniofacial cartilage - between its two extreme ends in both KO and missense homozygous larvae at 5 dpf. The phenotype was more severe in KO compared to missense larvae. An increase in the cartilage thickness of the Meckel’s was also observed in the KO model, but not in the missense larvae. Differences in phenotypic severity range of the cartilage likely reflect varying molecular consequences of the two variant models. The KO allele produces a truncated protein that may undergo nonsense mediated decay (NMD) or lose critical functional domains^41, 42^, whereas the missense variant may still produce a protein but with altered functional properties.

Further investigation of cartilage composition revealed variant-specific effects on type II collagen deposition. KO homozygous larvae exhibited increased thickness of the *col2a1*-positive cell layer, whereas homozygous missense larvae showed a subtle, but significant reduction in this layer. A range in phenotypic severity is also observed in ceratohyal cartilage, with the same thickness phenotype trend observed. Differences were also observed in *sox10*-positive chondrocytes, where the sphericity was different in *nbas* homozygous knockout larvae. Growth within the zebrafish jaw, primarily in the joints, is driven by cellular orientation and volume expansion^43^. Our data shows that development of the jaw appears to be disrupted through abnormal chondrocyte structures. Disorganised chondrocytes may limit the structural support to the developing cartilage and bone, potentially resulting in future bone fragility. Our findings indicate that *NBAS* variants produce a spectrum of cartilage and skeletal phenotypes rather than one uniform defect.

The NBAS protein has been implicated in NMD, where it cooperates with core NMD factors to regulate the expression of numerous RNA targets. Loss of NBAS function results in widespread transcriptional dysregulation^38^. Consequently, the cartilage and skeletal phenotypes observed in the KO model may arise not only from the disruption of NBAS-specific functions, but also from a broader dysregulation of other pathways regulated through NMD. By contrast, the missense protein is likely still produced and has a partial function to the wildtype protein, explaining the milder phenotypes observed in this model.

### Potential disease mechanisms underlying *nbas* cartilage and skeletal phenotypes

Another major role of NBAS is in retrograde transport between the Golgi apparatus and the endoplasmic reticulum (ER). The C-terminal domain of the protein is responsible for interacting with the p31-ZW10-RINT1 complex, which regulates COPI-mediated vesicle trafficking^44, 45^. Depletion of NBAS *in vitro* disrupts Golgi apparatus recycling processes^45^, highlighting the importance of these interactions for intracellular transport. Dysfunction of the Golgi apparatus is known to impair secretion during retrograde transport of proteins essential for cartilage and skeletal development, including collagens, which may also compromise post-translational modifications (PTMs) of these proteins, such as hydroxylation and glycosylation. Abnormal PTMs and protein secretion are well established contributors to skeletal and connective tissue disorders^46–49^. Collagen export from the ER also requires interactions between NBAS-associated protein complexes and TANGO1. Loss of these interactions in the absence of TANGO1 has been proposed to result in secretion of incorrectly folded collagen proteins, impacting their biological function^50^. Thus, a lack of NBAS may also result in improperly folded collagens, leading to developmental defects.

Our missense zebrafish model contains an amino acid substitution mutation in this C-terminal domain responsible for NBAS complex interactions with an arginine being replaced by a histidine. Arginine amino acid residues are known to generally form stronger interactions and bonds than histidine residues^51^. This amino acid substitution may reduce the stability of protein complexes without completely abolishing its formation and function. It may be the case that Golgi apparatus function and collagen processing may be partially retained in the missense model but the protein complexes are more unstable than when the NBAS protein is wildtype. Consequently, more subtle developmental abnormalities are found in our missense model.

In contrast, the KO variant model would be predicted to have more profound effects, which it does on overall skeletal and cartilage phenotypes. Two potential disease molecular mechanisms may account for the more severe phenotypes in this model. (1) Truncation of the protein eliminates the C-terminal domain completely, preventing the formation of essential protein complexes for collagen folding, PTMs and retrograde transport. (2) The KO transcript is targeted by NMD, resulting in the loss of NBAS protein expression and overall function. Both scenarios would impair Golgi apparatus-mediated recycling, collagen folding, and protein transport and secretion, and would be consistent with the more severe phenotype observed in KO homozygous larvae compared to missense homozygotes.

### Unexpected phenotypic outcomes in patient-specific compound heterozygous zebrafish model

The patient we previously identified with atypical OI was due to compound heterozygous variants in *NBAS* - with one allele being a premature termination codon nonsense variant, and the missense arginine to histidine substitution on the other. We were able to model this specific patient genotype with our zebrafish. It was expected that an intermediate or all-round similar phenotype would be observed in a compound heterozygous model compared to the separate allelic models, as has been seen in compound heterozygous zebrafish models of human hypothyroidism^52^. Surprisingly, the compound heterozygous larvae exhibited somewhat opposing phenotypes in certain phenotypic parameters. Increased operculum length ratio and Meckel’s cartilage arc length were seen at 5 dpf, while mineralisation and other phenotypic parameters were unaffected by the compound heterozygous genotype. These are in contrast with the shorter morphometric measurements observed with our other genotype models.

The molecular basis of this unexpected compound heterozygous phenotype remains unclear. One possibility is intragenic complementation, where proteins derived from different mutant alleles interact to form a partially functional complex with greater activity than either mutant protein found alone^53^. This mechanism could explain the apparent reversal of operculum phenotypes. Alternatively, compound heterozygosity may have a neomorphic effect on the produced protein, resulting in a gain-of-novel-function^54^. This would produce a protein with different functions to the wildtype and mutant proteins alone, resulting in the differing phenotypes observed. Our current data suggests that the compound heterozygous model has phenotypes which are partially rescued, but that this rescue may extend beyond normal physiological limits. Additional biochemical and functional studies of the protein will be required to determine the exact molecular mechanisms involved and the functional impact of compound heterozygosity.

### Conclusion and future perspectives

Collectively, these findings demonstrate that *NBAS* variants produce a complex spectrum of skeletal and cartilage phenotypes that are dependent on the mutation type. This study establishes zebrafish models for investigating *NBAS*-associated atypical OI, and provides insights into the developmental consequences of truncating, missense, and compound heterozygous variants. These models offer a valuable platform for future work delineating molecular mechanisms of each variant, but also future studies for therapeutics and development of treatment strategies for this rare, but life changing, genetic disorder.

## ACKNOWLEDGEMENTS

We would like to thank the patients and their families for their participation in this study. We thank Henry Roehl, Heba Ismail, Stephen Renshaw, Iwan Evans, Phil Elks, Simon Johnston, and Gillian Tomlinson for their feedback and critique of our work. We would also like to thank all the Biological Services Aquaria Team for fish care and practical assistance. We are grateful to Sheffield Children’s Hospital Charity for funding. FvE was supported by BBSRC grant BB/R015457/1. MB received support from MRC Fellowship (MR/V037307/1) from September 2021 to August 2024.

## AUTHOR CONTRIBUTIONS

MB conceived the study. MB, DB, and ZM supervised experiments and analyses. DB, MB and ZM wrote the manuscript with input from all authors (SS, KM, ASA, AA, MZ, NS, SB, CL, FvE, HR). Clinical data and literature review was undertaken by KM and ASA. Zebrafish models generated in-house were done by FvE, CL and MB. Zebrafish experiments and analyses were performed by DB, SS, ZM, AA, MZ, NS, CL, and FvE.

## SUPPLEMENTAL FIGURE LEGENDS

**Suppl. Figure 1.**
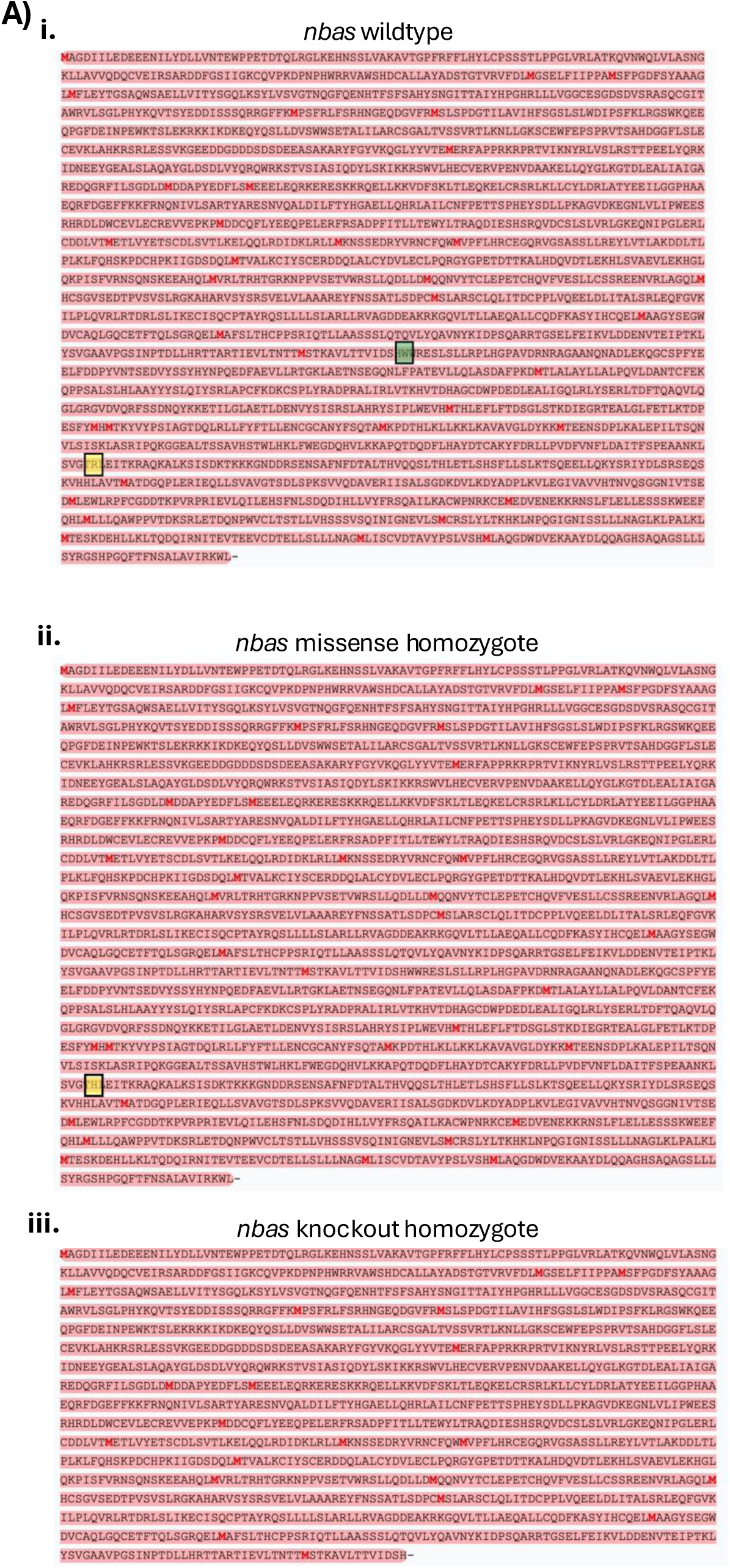
Impact of *nbas* mutations on amino acid sequence. **A.** Amino acid sequence for Nbas zebrafish protein predicted in **i)** wildtype, **ii)** missense homozygotes, and **iii)** knockout homozygotes.

**Suppl. Figure 2.**
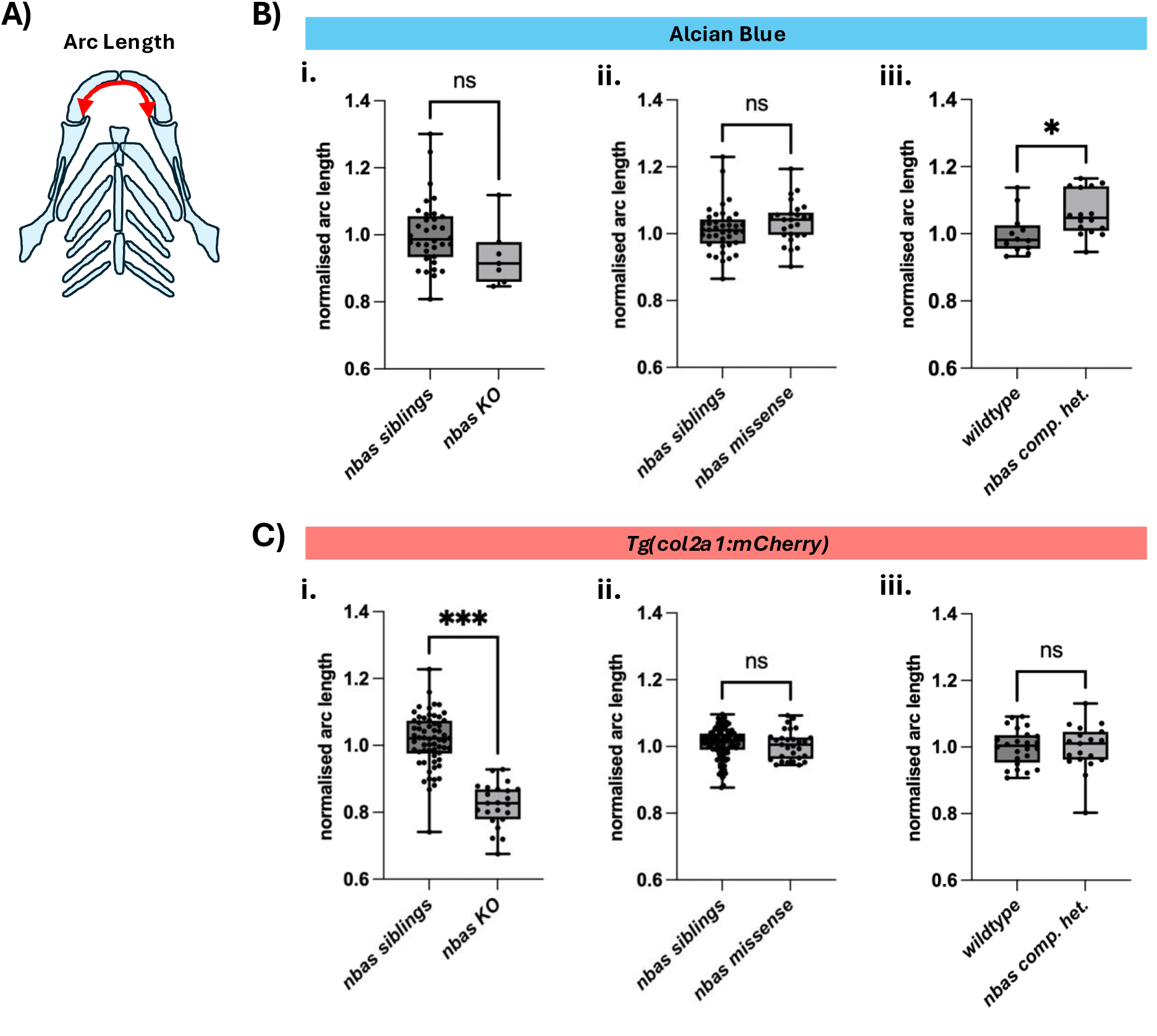
Arc length measurements of Meckel’s craniofacial cartilage. **A.** Schematic showing how arc length of the Meckel’s cartilage is taken, highlighted by the red arrow. **B.** Quantification of Meckel’s cartilage arc length from Alcian blue stainings at 5 dpf for **i)** knockout, **ii)** missense, and **iii)** compound heterozygous larvae. An unpaired t test was used for statistical analyses. **C.** Arc length measurement quantifications from *Tg(col2a1:mCherry)* fluorescent images at 5 dpf for **i)** knockout, **ii)** missense, and **iii)** compound heterozygous larvae. Either an unpaired t test or a Mann-Whitney test were used for statistical analyses. ns, not significant. * *p* < 0.033. *** *p* < 0.001.

**Suppl. Figure 3.**
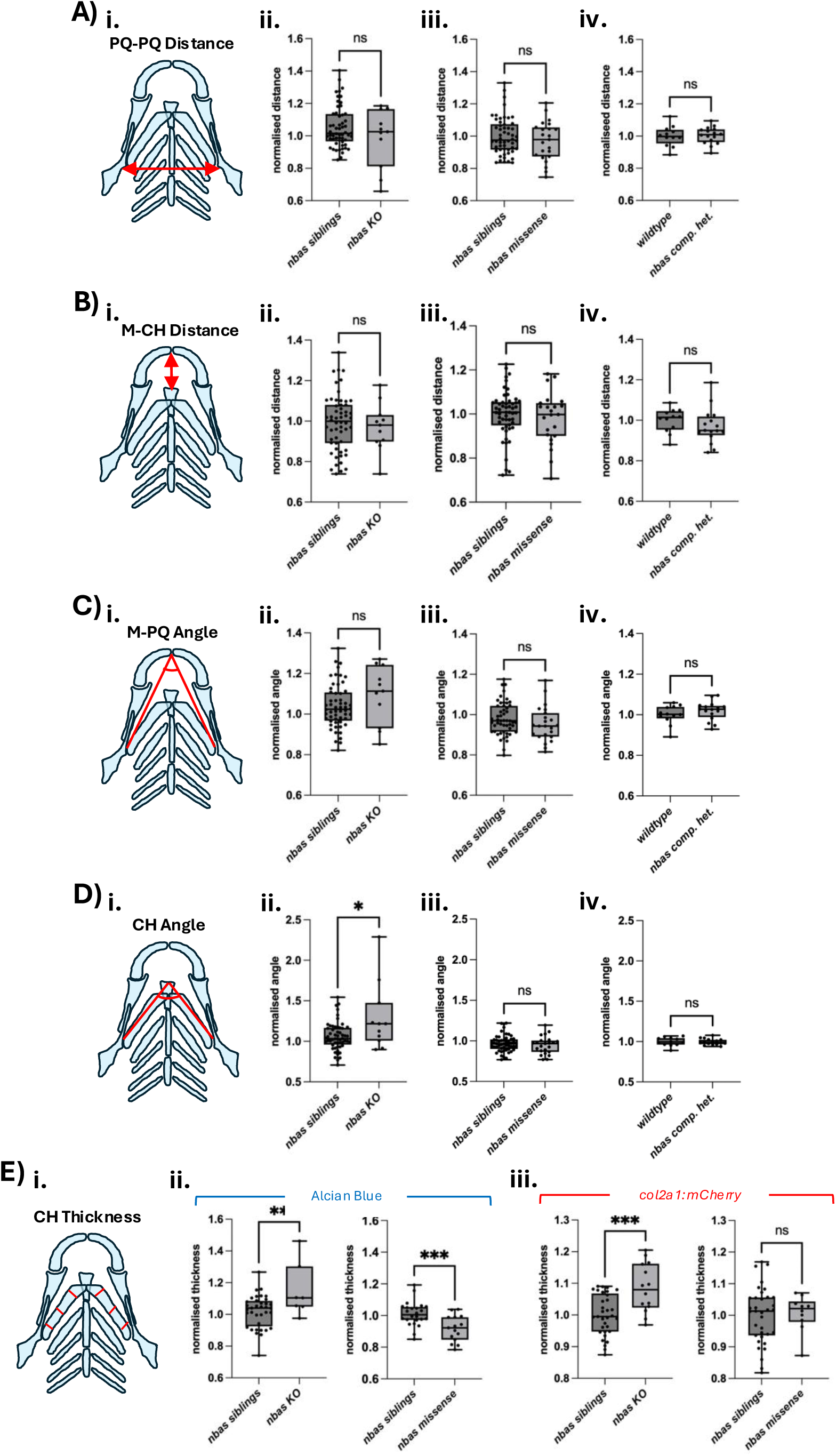
Quantification of additional craniofacial cartilage parameters. **A.** Analysis of palatoquadrate-palatoquadrate (PQ-PQ) distance. **i)** Schematic showing PQ-PQ distance measurement with red arrow. Quantification of PQ-PQ distance at 5 dpf for **ii)** knockout, **iii)** missense, and **iv)** compound heterozygous larvae. Either an unpaired t test or a Mann-Whitney test were used for statistical analyses. **B.** Analysis of Meckel’s-ceratohyal (M-CH) distance. **i)** Schematic showing M-CH distance measurement with red arrow. Quantification of M-CH distance at 5 dpf for **ii)** knockout, **iii)** missense, and **iv)** compound heterozygous larvae. Unpaired t tests were used for statistical analyses. **C.** Analysis of Meckel’s-palatoquadrate (M-PQ) angle. **i)** Schematic showing M-PQ angle measurement. Quantification of M-PQ angle at 5 dpf for **ii)** knockout, **iii)** missense, and **iv)** compound heterozygous larvae. Unpaired t tests were used for statistical analyses. **D.** Analysis of ceratohyal (CH) angle. **i)** Schematic showing CH angle measurement. Quantification of CH angle at 5 dpf for **ii)** knockout, **iii)** missense, and **iv)** compound heterozygous larvae. Either an unpaired t test or a Mann-Whitney test were used for statistical analyses. **E.** Analysis of ceratohyal (CH) cartilage thickness. **i)** Schematic showing CH cartilage thickness measurements done by averaging six measurements taken from across the cartilage. Quantification of CH normalised thickness layers for knockout and missense 5 dpf larvae in **ii)** Alcian blue images and **iii)** *Tg(col2a1:mCherry)* fluorescent images. Unpaired t tests were used for statistical analyses. ns, not significant. * *p* < 0.033. ** *p* < 0.002. *** *p* < 0.001.

